# Isolation and characterisation of watermelon (*Citrullus lanatus*) extracellular vesicles and their cargo

**DOI:** 10.1101/791111

**Authors:** Kate Timms, Beth Holder, Anil Day, John McLaughlin, Melissa Westwood, Karen Forbes

## Abstract

Extracellular vesicles (EVs) facilitate cell-cell communication in animals and are integral to many physiological and pathological processes. Evidence for the presence and function of EVs in plants is limited. Here, we report that EVs derived from watermelon fruit mesocarp are of similar size and morphology to the animal EV subtype known as exosomes. Analysis of EV constituents revealed that watermelon EVs are negative for endoplasmic reticulum markers, and that the miRNA and protein profiles differ from that of watermelon mesocarp cells, suggesting that these EVs are actively synthesised and are not merely cellular debris. Furthermore, we report a panel of proteins found in in watermelon EVs as well as the published proteomes of grape, grapefruit, lemon and *Arabidopsis thaliana* EVs that are novel potential plant EV markers. Bioinformatic analyses suggest that plastids and multivesicular bodies are likely sites of biogenesis for EVs from watermelon and other plants. Predicted functional roles of watermelon EVs include development and metabolism, with several of their cargo molecules likely to be key in regulation of fruit development and ripening. Further understanding of how EVs may contribute to these processes would improve understanding of plant cell-cell communication and could aid in the harnessing of plant EVs for greater temporal control of crop development/ripening for the agricultural and retail industries.

## Main

An ever-growing literature emphasises the critical role played by extracellular vesicles (EVs) in mediating cell-cell communication, both in health and disease, in the bacterial, fungal and animal kingdoms^1,2^. These EVs travel in the extracellular environment before internalisation by recipient cells, delivering their cargo and altering cell function^3-5^. Whilst EV-like particles have been observed in plants ^6-16^, their identity as true EVs (i.e., vesicles that are actively secreted, rather than mere cellular fragments) is yet to be confirmed. The term ‘EVs’ encapsulates several classes of cellular-derived vesicles with varying size, composition and function. One of most studied classes of mammalian EVs are exosomes, which are derived from membrane invagination in the late endosomes known as multivesicular bodies (MVBs)^17,18^. Exosomes are generally considered to be 50-150nm EVs that contain a range of cargo molecules, including proteins, messenger RNAs (mRNAs) and microRNAs (miRNAs), which are actively sorted into exosomes during exosome formation, resulting in a differing cargo profile to that of the cell of origin^5,19-24^. Differing cell and EV cargo demonstrates that EVs are purposeful and energy demanding cell-cell signals, not just passive cellular products. Microvesicles are larger (100nm-1µm) EVs which bud directly from the plasma membrane^25^, meaning that their contents more closely resemble that of the donor cell. The third major EV subtype is the 1-2µm apoptotic bodies, which bleb off the plasma membrane in apoptotic cells^26,27^ and can contain whole organelles or nuclear fragments^28^.

In plants, MVB fusion with the plasma membrane at sites of infection has been observed using electron microscopy (EM), and is thought to be involved in the response to pathogens^29^. Plant EVs have been shown to contain small RNAs^30^, such as miRNAs, which are suggested to suppress virulence genes in fungal pathogens^31,32^. Despite these more recent studies, majority of the studies reporting EV-like particles in plants^6-16^ have focused on dietary ingestion of plant EVs by animals, meaning that the particles described in such studies have been characterised largely in the context of their impact on mammalian cells, rather than the plant they originated from^8,11,13^. Initial reports on the presence of membrane phospholipids in plant extracellular fluid^33,34^ were followed by a study aiming to determine whether these lipids were within EVs ^15^. Using the classical differential ultracentrifugation method of EV isolation followed by EM, EV-like particles were visualised in pellets from apoplastic fluid at both 40,000xg and 100,000xg^15,35^. In keeping with their potential identity as EVs, these particles were found to contain at least a limited protein cargo. Further characterisation of the components, biogenesis and function of plant EVs has yet to be performed. Determining the origin and function of plant EVs is essential to understanding plant cell-to-cell communication, not only to increase fundamental knowledge of plant physiology, but in understanding how these processes may be beneficially manipulated. Watermelon fruits (*Citrullus lanatus*) contain an abundance of extracellular fluid and have, at ripeness, a delicate mesocarp, allowing for the isolation of large quantities of extracellular fluid without risk of disrupting cells and creating cellular fragments which could erroneously be identified as EVs. As such, watermelon fruit is an ideal model for plant EV characterisation. Here, we show that watermelon fruit contains EVs – likely originating from plastids and MVB – which differ in miRNA and protein profiles to mesocarp cells. Furthermore, we identify a panel of potential plant-specific EV markers and predict that watermelon EVs have likely roles in sugar metabolism and fruit development/ripening.

## Results

### Characterisation of extracellular vesicles from watermelon fruit mesocarp

Sequential ultracentrifugation was used to isolate EVs from watermelon mesocarp. Parallel isolation from culture medium conditioned by human epithelial cells was performed to validate the isolation procedure. The presence of authenticated human EV markers^36-39^ (ALIX^+ve^/CD63^+ve^/CANX^-ve^; Figure 1a) in the EVs from human epithelial cell conditioned media confirmed the suitability of our approach. Nanoparticle tracking analysis and transmission electron microscopy revealed that watermelon and human EVs have a similar morphology (Figure 1b) and size (117nm ± 14.61 Vs 129nm ± 28 respectively, p=0.68; Figure 1b). However, it should be noted that the watermelon Evs isolated here appear to consist of 2 subsets, one with a peak around 40-60nm in diameter and one around 150nm, potentially reflecting two distinct subtypes of EVs. The average yield of watermelon EVs was 1.85×10^10^ ± 1.63×10^10^ EVs/ml of watermelon juice. On average, the DNA yield was 2.12×10^12^ ± 9.57×10^11^, the RNA yield was 7.64×10^8^ ± 9.31×10^8^ particles/ng RNA and the protein yield was 1.94×10^9^ ± 7.14×10^8^ particles/μg protein. Relative to the starting volume of juice, the EV DNA yield was 0.38 ± 0.11 μg/ml, the EV RNA yield was 3.45 ± 1.70 μg/ml and the EV protein yield was 96.79 ± 32.6 ng/ml.

**Figure 1.**
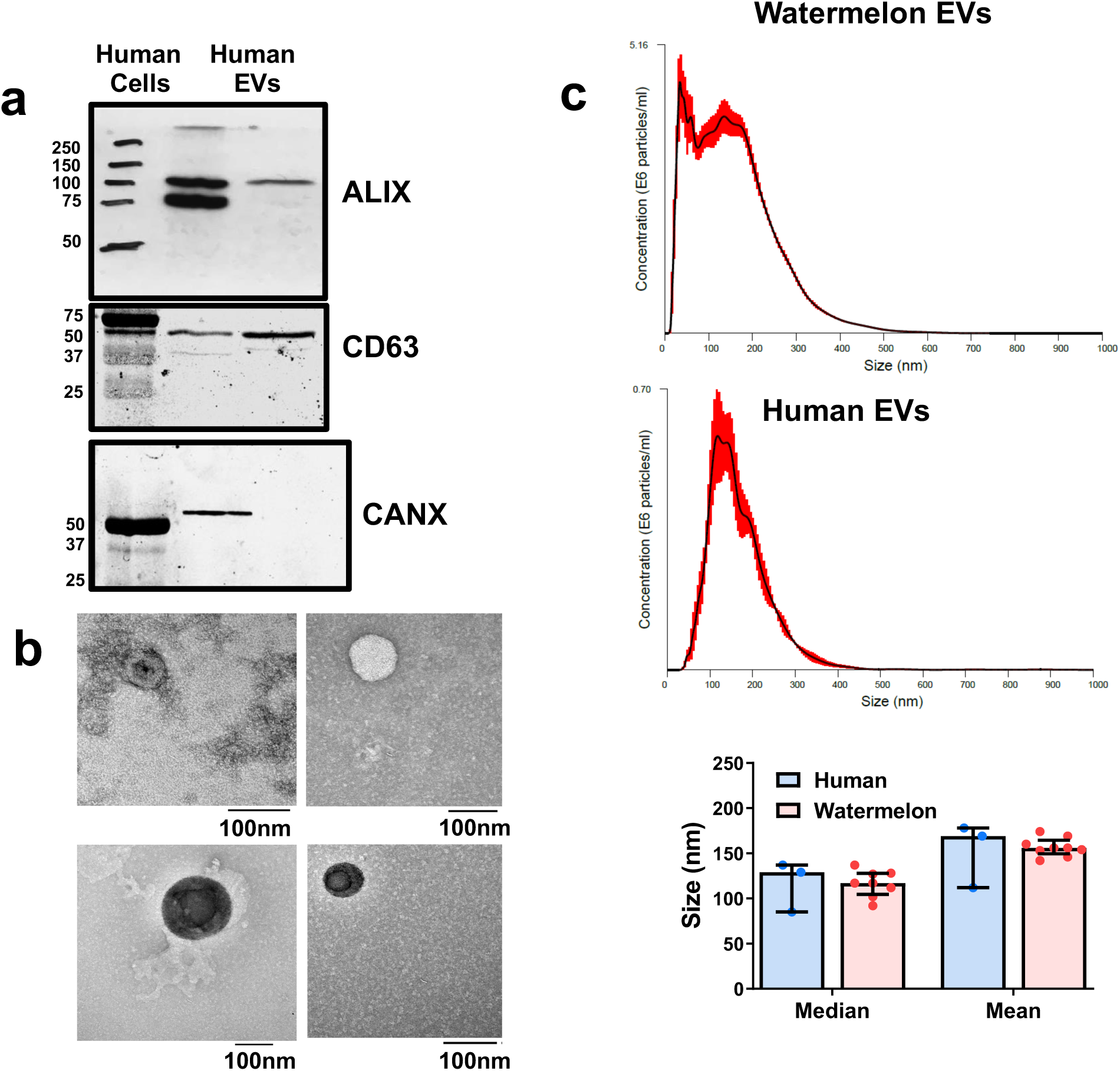
Watermelon extracellular vesicles (EVs) resemble human exosomes. **a**, Western blot analysis of positive (ALIX and CD63) and negative (CANX) markers for exosomes in EVs isolated by differential ultracentrifugation from the conditioned media of human Caco-2 cells. **b**, Watermelon EVs display vesicular morphology reminiscent of human exosomes when analysed using transmission electron microscopy. EVs show characteristic cup-shaped morphology. **c**, Size comparison of watermelon and human derived EVs. Particles isolated by ultracentrifugation from watermelon mesocarp and human Caco-2 cell conditioned culture media were subjected to nanoparticle tracking analysis to determine size. Representative size distribution graph for watermelon EVs (n=8) and human EVs (n=3) for both mean (p= 0.6364) and median size (p= 0.6788) are shown. Data are shown as median ± IQR. (n=3-8; Mann-Whitney test).

### Protein and miRNA profiling of watermelon EVs

Next, to rule out the possibility that the isolated watermelon EVs were merely products of cellular disruption or breakdown, we compared their protein and miRNA content to that of their parent mesocarp cells. The full list of proteins identified (1838) in watermelon EVs and cells is available in Supplementary table S1. 28.13% of proteins were enriched and 34.98% were depleted by >1.5-fold in EVs compared to parent cells. Of note, a number of cell compartment markers for the endoplasmic reticulum (ER), mitochondria, nucleus and chloroplast, including established negative markers of mammalian EVs such as ER-localised calnexin and nuclear-localised histone 2B, are depleted in EVs compared to cells (Table 1). Principal component analysis and hierarchical clustering analysis indicated that the profile of proteins in watermelon mesocarp EVs differed from that of cells, though the profile varied between different watermelons (Figure 2a-b). Further analysis of watermelon EV content was performed by profiling the miRNAs. This demonstrated that 25.00% and 35.71% were >1.5-fold enriched and depleted respectively in EVs compared to mesocarp cells (Figure 2c-e; full results in Supplementary table 2). Principal component analysis confirmed that EVs have a distinct profile of miRNAs compared to their parent cells (Figure 2c).

**Table 1:** Depletion of cell compartment markers in watermelon extracellular vesicles (EVs) in comparison to watermelon cells.

| **Cellular Compartment** | **Accession** | **Protein Name** | **Relative abundance in Watermelon Cells** | **Relative abundance in Watermelon EVs** |
| --- | --- | --- | --- | --- |
| Nucleus | Cla019920 | Histone H2A | 1.33 | 0.34 |
|  | Cla001861 | Histone H2B | 2.334 | 0.34 |
|  | Cla015219 | Histone H3 expressed | 1.34 | 0 |
|  | Cla005035 | Histone H4 | 4.34 | 0.67 |
| Endoplasmic Reticulum | Cla002648 | Calnexin | 7.34 | 0.33 |
|  | Cla013928 | Protein disulfide isomerase | 11.83 | 7.33 |
|  | Cla005127 | Cytochrome P450 | 2.67 | 0.00 |
| Mitochondria | Cla000734 | Cytochrome c1 | 4 | 0 |
|  | Cla016943 | Cytochrome C-2 | 1.00 | 0.34 |
|  | Cla022586 | Mitochondrial ADP/ATP carrier protein 3 | 8.34 | 1.67 |
|  | Cla002230 | Mitochondrial ADP/ATP carrier protein 1 | 2.67 | 0 |
| Chloroplast | Cla010898 | Phytoene desaturase | 7 | 2.67 |
|  | Cla006309 | Sulfite reductase | 5 | 0.34 |
|  | Cla008831 | Plastid lipid-associated protein 3, chloroplastic | 1.67 | 0.67 |
|  | Cla000424 | Chloroplast outer envelope protein 37 | 2 | 1.67 |
|  | Cla020214 | Zeaxanthin epoxidase chloroplastic | 1 | 0 |
|  | Cla008113 | Tic110 family transporter chloroplast inner envelope protein | 4.33 | 1 |

**Figure 2:**
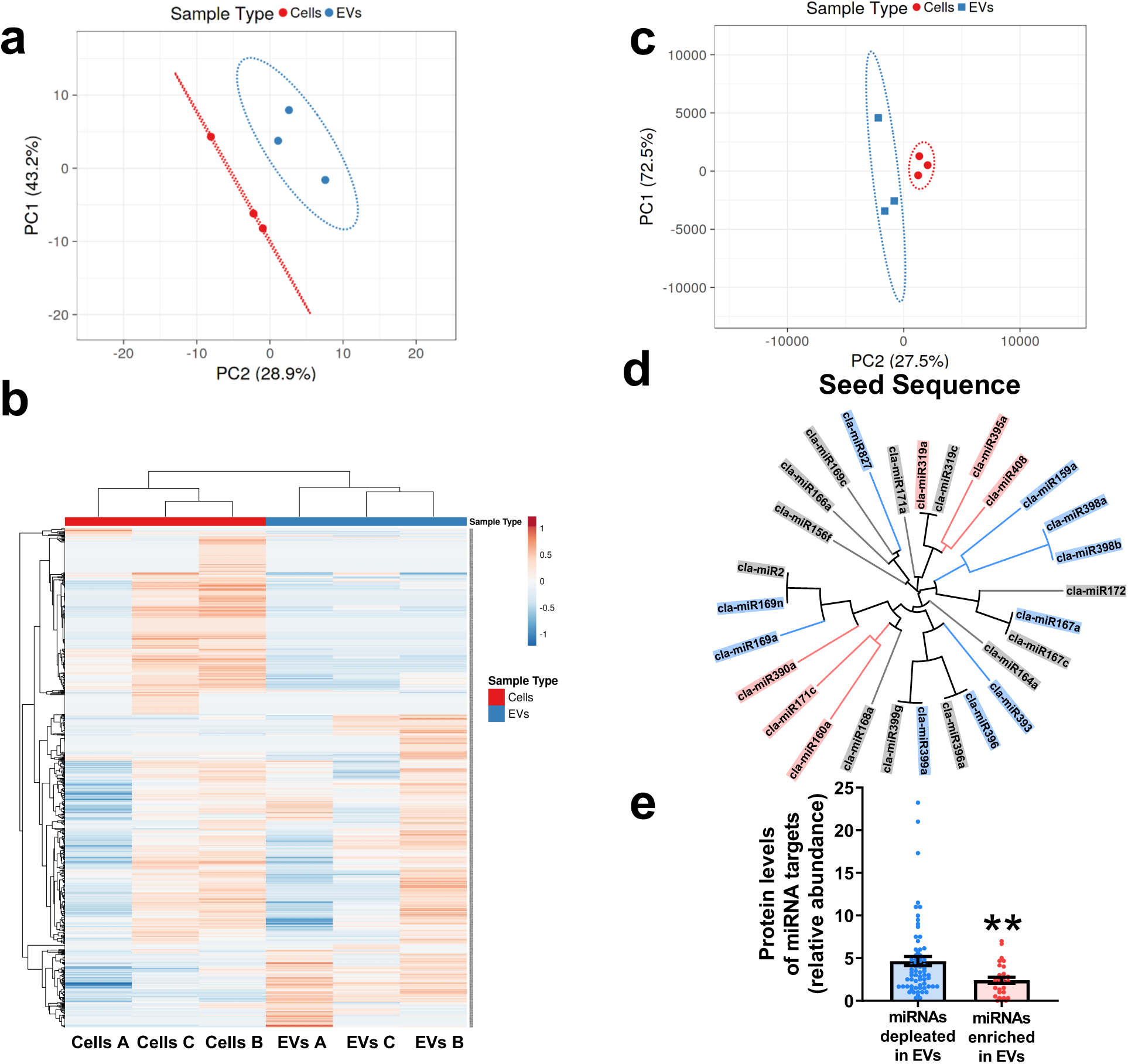
Characterisation of the watermelon extracellular vesicle (EV) proteins and miRNAs. **a-b** The proteome of watermelon mesocarp EVs differs from that of cells. Watermelon protein profiles (n=3) were analysed using principal component analysis (**a**; PCA; log10(relative abundance)) or hierarchical clustering (**b**). **c-e**, miRNAs with a 1.5-fold higher/lower level in EVs than in cells were designated as enriched/depleted (n=3). **c**, PCA of miRNA levels in mesocarp cells and EVs (log10(ΔΔCt)). **d**, miRNA relationship based upon 8-base seed sequences, coloured for enrichment (red), depletion (blue) or neutral (grey) localisation in EVs with respect to cells. **e**, Protein levels of mRNAs targeted by miRNAs are enriched (n=29) or depleted (n=67) in EVs. Data are shown as mean ± SEM (Mann-Whitney test; p= 0.0045; ** = p<0.01).

Together these data suggest that the EVs isolated in this study are not merely fragments of organelles produced during cellular breakdown or EV isolation. The enrichment of specific miRNAs in watermelon EVs compared to cells suggests active sorting of miRNAs in watermelon. In mammals, miRNAs are actively sorted into EVs, in a sequence-dependent manner, by hnRNPA2B1^23^. To determine whether the same may be true of plant EVs, a BLAST search for hnRNPA2B1 was conducted. No sufficiently orthologous protein was found in the Viridiplantae kingdom. Furthermore, analysis of the relationship between miRNA sequence, sequence motif or secondary structure and enrichment/depletion in watermelon EVs (data not shown) revealed no obvious association. However, when the miRNA seed sequence was considered in isolation, it did appear that some related seed sequences were similarly regulated (Figure 2d). As the seed sequence of miRNAs is the most important region for miRNA-mRNA binding, and mRNA abundance is inversely linked to miRNA sorting EVs in animals^40^, we hypothesised the relationship between seed sequence and EV enrichment may relate to the abundance of target mRNAs within watermelon cells. Whilst the levels of mRNAs in watermelon cells are unknown, our analysis of the proteins present in watermelon cells and EVs can be used as a proxy measure of the abundance of their mRNAs; our data show that the levels of the cellular proteins that are the targets of miRNAs enriched in EVs were lower than those that are the targets of miRNAs depleted in EVs (Figure 2e; p<0.01). A full list of watermelon EV miRNA targets is given in Supplementary Table 3.

### Identification of potential plant EV markers

There is a growing consensus on the use of several lipid-associated/membrane-spanning and cytosolic proteins as suitable markers of mammalian EVs^39^. Likewise, there is a proposed range of ‘negative markers to distinguish human EVs, particularly the MVB derived exosomes, from cellular debris^39^. We also demonstrated the depletion of some of these ‘negative’ markers, such as calnexin, in watermelon EVs compared to watermelon cells (Table 1).

Currently, there are no widely recognised positive markers of plant EVs and there are no sufficient homologs of human markers present in the watermelon proteome, suggesting mechanisms of biosynthesis may differ. To discover potential novel marker(s) of plant EVs, the proteomes of EVs from watermelon and EV-like particles previously isolated from grape^11^, grapefruit^13^, lemon^14^ and *Arabidopsis thaliana*^6^ were compared (Table 2). One protein, enolase, is present in the EVs of all 5 species and, interestingly, is the second most commonly reported protein in human EVs^41^. Of the 15 remaining potential markers with a human ortholog, all have been identified in human EVs and 11 are within the top 250 most commonly identified human EV proteins^42^.

**Table 2:**
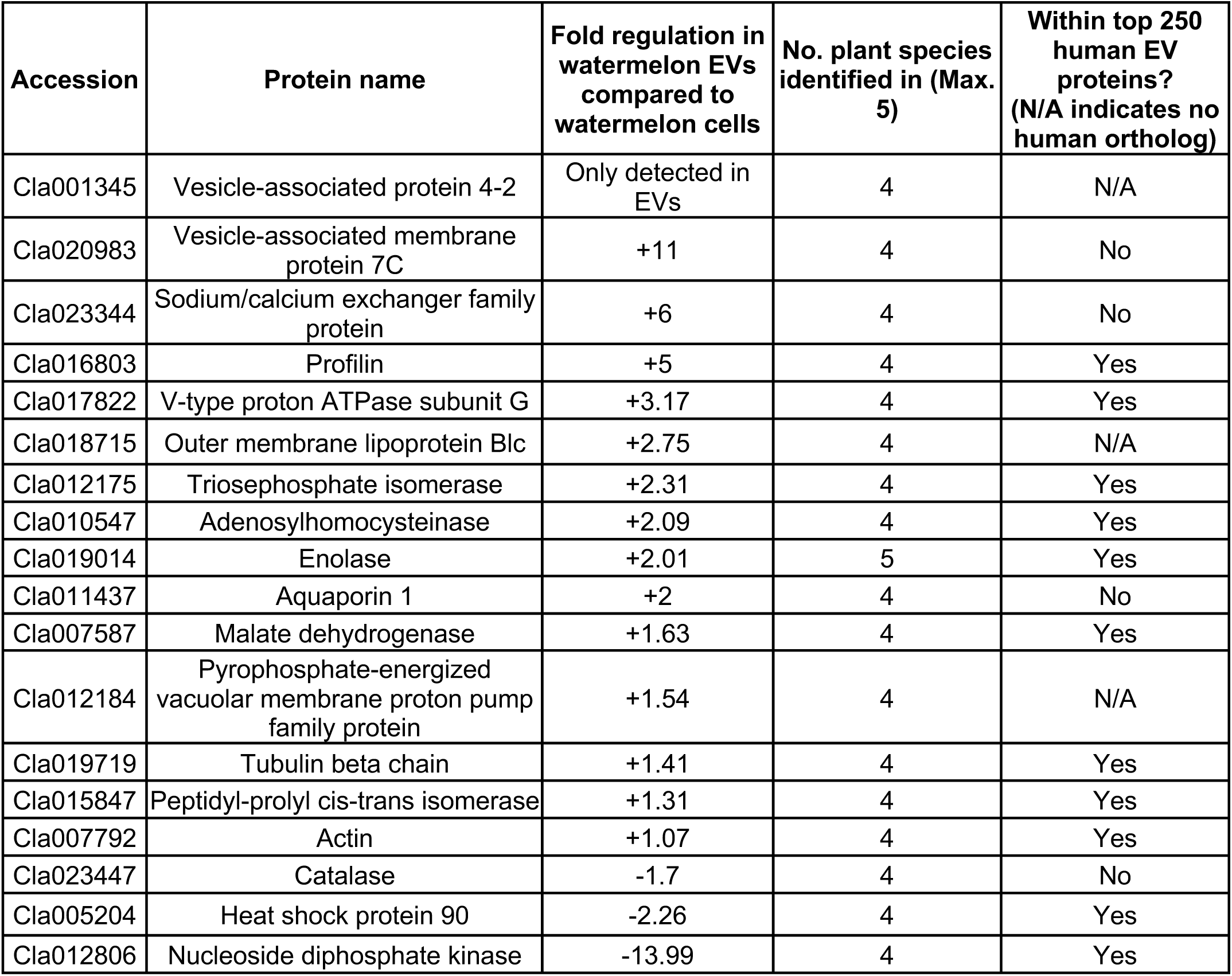
Candidate plant extracellular vesicle markers.

### Plastids and multivesicular bodies are likely sites of watermelon extracellular vesicle biogenesis

We next interrogated the EV protein profile using gene ontology (GO) enrichment analysis, reasoning that such information might help to identify the site of EV biogenesis. This indicated that the majority of the EV proteins are from either the cytoplasm or the plastid and its derivatives (Figure 3a). The plastid is a known site of vesicles^43-49^ approximating the size of the smaller peak identified in nanoparticle tracking analysis of watermelon EVs (Figure 1c), making the plastid a credible candidate site for plant EV biogenesis. Further prediction based on cellular localisation signals and comparison of the watermelon EV proteome with previously published data^50^ supports this hypothesis, with 73.58% of watermelon EV proteins predicted/observed to be within plastids (Figure 3b). The same high percentage of plastid proteins was seen in other plant species for which data was available.

**Figure 3:**
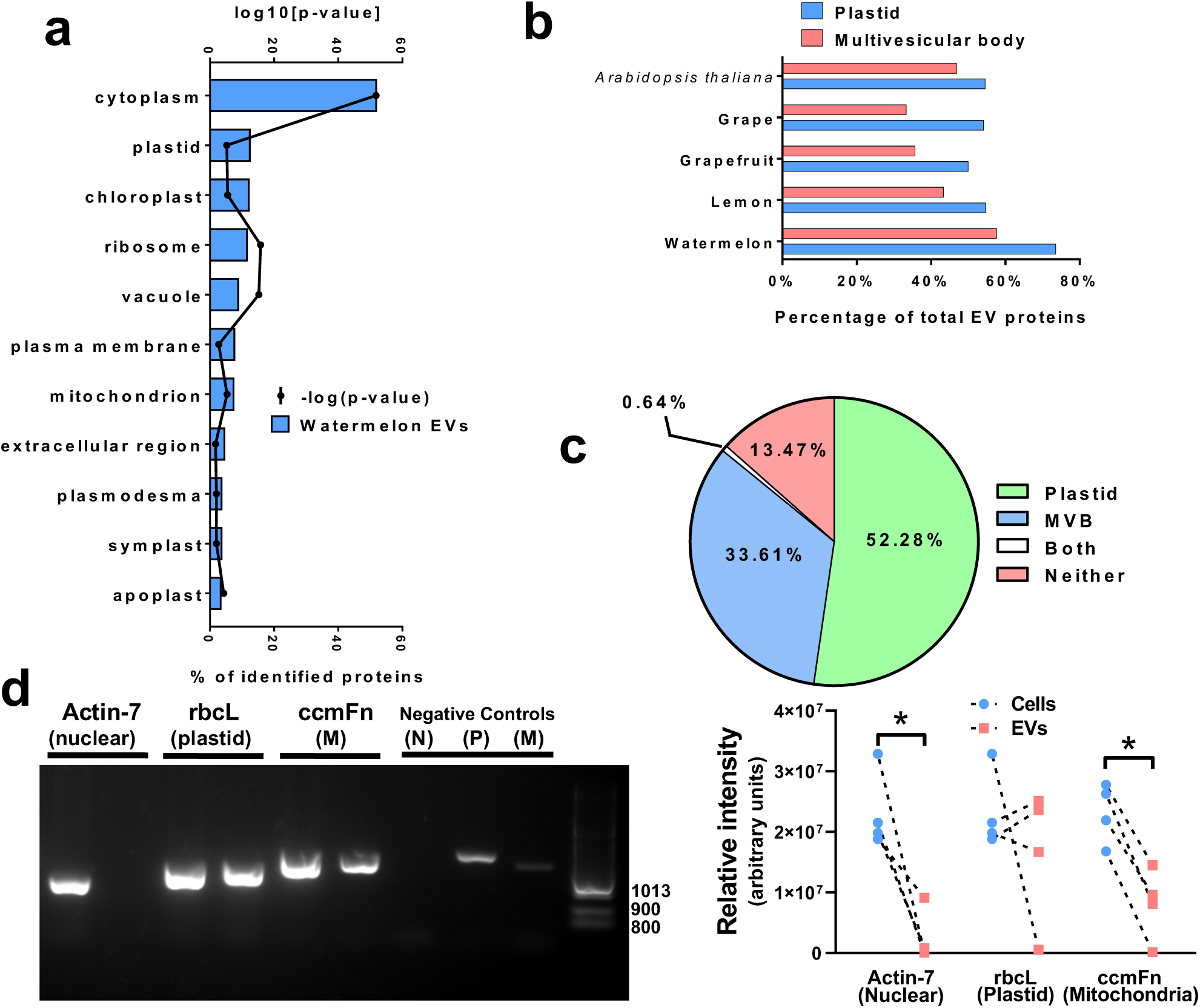
Watermelon EVs are likely derived from plastids and multivesicular bodies. **a**, GO enrichment of watermelon EV protein localisation (Bonferroni p-value correction). **b**, Comparison of the percentage of multivesicular body (MVB) and plastid proteins present in EVs from watermelon and those previously published for EVs in lemon^14^, grapefruit^13^, grape^11^ and *Arabidopsis thaliana*^6^ were aligned to the proteomes of watermelon chromoplasts^50^ and Arabidopsis thaliana multivesicular bodies^51^. **c**, A comparison of the percentage of the total watermelon EV protein mass predicted and/or reported to reside cellularly within plastids, MVB, both or neither organelle. **d**, Relative quantification of DNA of nuclear (N; actin-7; p=0.0286), plastid (P; rbcL; p= 0.6857) and mitochondrial (M; ccmFn; p= 0.0286) origin in EVs compared to cells (n=4). Data are shown as median (Mann-Whitney test; * = p<0.05).

Furthermore, the yellow-orange colouration of watermelon EV pellets suggests the presence of lycopene, which is produced and stored within watermelon chromoplasts. Interestingly, our analysis of the published proteomes of EV-like particles isolated from grape^11^, grapefruit^13^, lemon^14^ *and Arabidopsis thaliana*^6^ suggests that plastids might be a common site of EV synthesis in plants (Figure 3b).

Animal exosomes originate from MVB. As neither GO enrichment nor predictive localisation based upon sequence are equipped for the assessment of MVB proteins, BLAST was used to align the proteome of EVs from watermelon, grape, grapefruit, lemon and *Arabidopsis thaliana* to that of *Arabidopsis thaliana* MVB^51^. This revealed that the majority of the non-plastid EV proteins are present in plant MVB, with some proteins being present in both organelles (Figure 3b). When protein abundance is considered, rather than the number of individual proteins, 52.28% of all protein mass in watermelon EVs is from proteins which reside solely in plastids, whilst proteins residing solely in MVB account for just 33.61% of the total protein mass in watermelon EVs (Figure 3c), suggesting that EVs of potential plastid origin may be more abundant than those of MVB origin.

Proteins are not the only plastid component present in EVs. Whilst DNA of nuclear (actin-7) and mitochondrial (ORF204) origin are depleted in EVs compared to cells (Figure 3d; p<0.05), the level of plastid DNA (rbcL) in cells and EVs was similar. The level of plastid DNA present in EVs varied between watermelons. Nonetheless, these data further support a plastid origin for at least a subset of watermelon EVs.

### Predicted roles for watermelon extracellular vesicles in fruit development/ripening

Next, we turned to predicting the potential roles of watermelon EVs by examining GO cellular process and pathway enrichment for EV proteins and miRNA targets. The targets of miRNAs that are enriched in cells appear to be concerned mainly with the various processes of cellular metabolism (Figure 4; blue), whereas the targets of miRNAs enriched in watermelon EVs are primarily involved in processes of development and reproduction (Figure 4; red). Analysis of the proteins enriched in watermelon cells and EVs suggests that they have broadly similar roles in both compartments (Figure 5a). However, whilst the processes and pathways of proteins enriched in cells cover a wide range of cellular metabolic processes, those for proteins enriched in EVs focus on nucleic acid metabolism, suggesting a role in gene expression. When considered in isolation, those watermelon EV proteins that are predicted to usually localise to the MVB or the plastid are also mostly involved in metabolism (Figure 5b). The second highest enriched biological process theme in MVB is gene expression (11.1%), with nucleic acid metabolism also featuring in the metabolism processes. However, in plastids, the second highest enriched biological process theme is response to reactive oxygen species (4.61%), followed closely by cell signalling (4.15%), and gene expression does not feature. These data suggest distinct roles for the proposed MVB-derived and plastid-derived EV subtypes.

**Figure 4:**
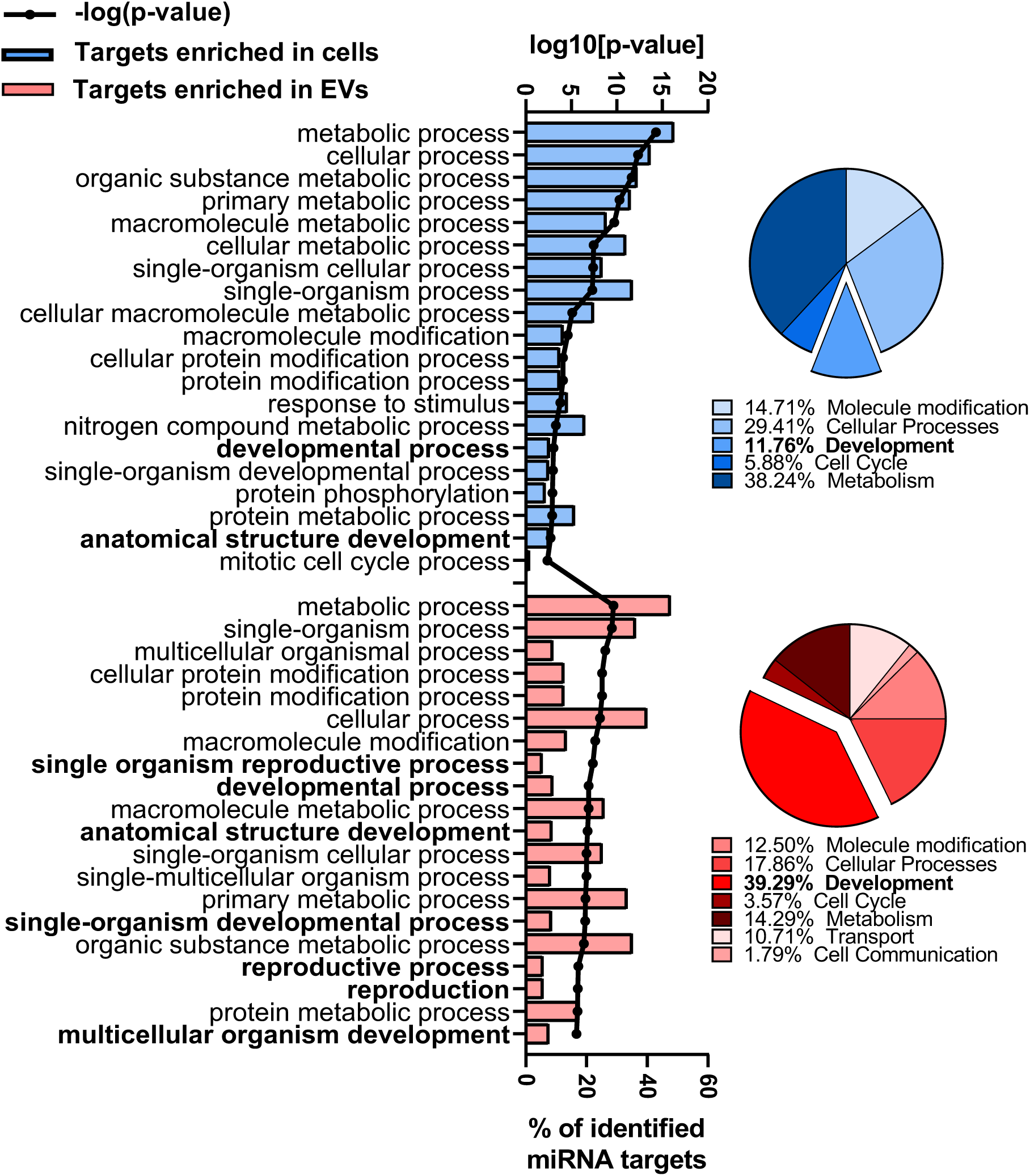
A potential role for watermelon EVs miRNAs in fruit development/ripening. Functions of miRNAs (via their predicted mRNA targets) which are enriched in watermelon cells (blue) or EVs (red) were predicted through GO biological process enrichment analysis. Top biological processes based upon p-value are given in the bar chart and all processes grouped by theme given in the pie charts (development emphasised). p-value correction for multiple testing was preformed using the Bonferroni method.

**Figure 5:**
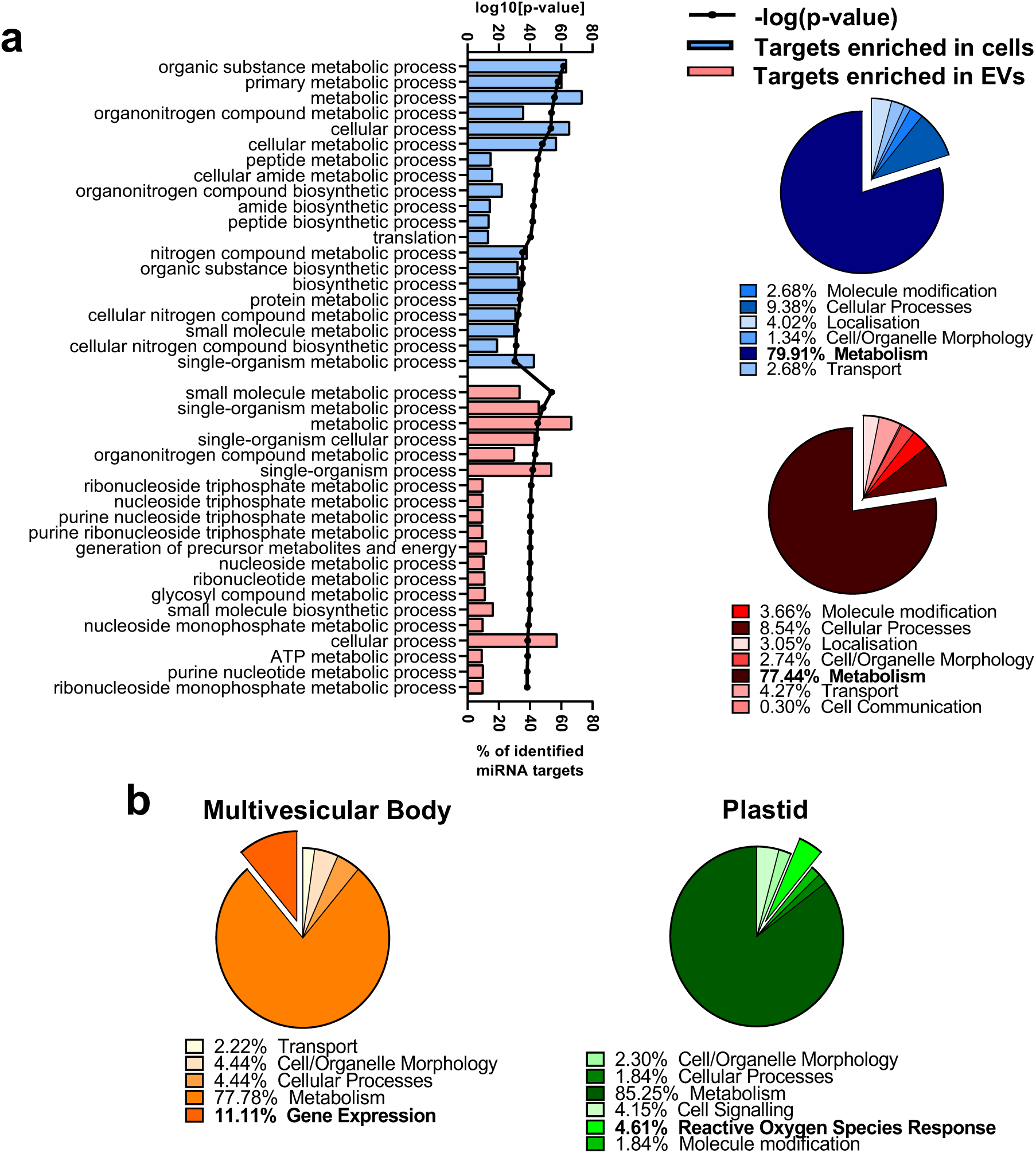
A potential role for watermelon EV proteins in metabolism, gene expression and reactive oxygen species degradation. **a**, Functions of proteins which are enriched in watermelon cells (blue) or EVs (red) were predicted through GO biological process enrichment analysis (Bonferroni p-value correction). The most significantly enriched biological processes are given in the bar chart and theme-grouped processes in the pie charts (metabolism emphasised). **b**, Biological process enrichment for watermelon EV proteins predicted to localise to the multivesicular body or plastid. The top biological process theme following from metabolism is emphasised.

As development/reproduction, metabolism and control of reactive oxygen species are all important for fruit production and ripening, we hypothesised that watermelon EVs could have a role in these processes. We therefore analysed an open access dataset^52^ of the transcripts expressed in the fruit flesh of the LSW177 variety of watermelon on days 10-34 after pollination, predicting that the mRNA targets of miRNA found in watermelon EVs would be downregulated over time if EVs truly influence fruit development/ripening. The data presented in Figure 6 support this hypothesis as the levels of watermelon EV miRNA targets decrease across the timescale of fruit development and ripening, reaching significantly lower expression than the average of all watermelon transcripts at day 34 (p<0.0001) when the fruit is considered ripe. Moreover, the transcripts for proteins present within watermelon EVs were enriched compared to all watermelon transcripts across this entire period.

**Figure 6:**
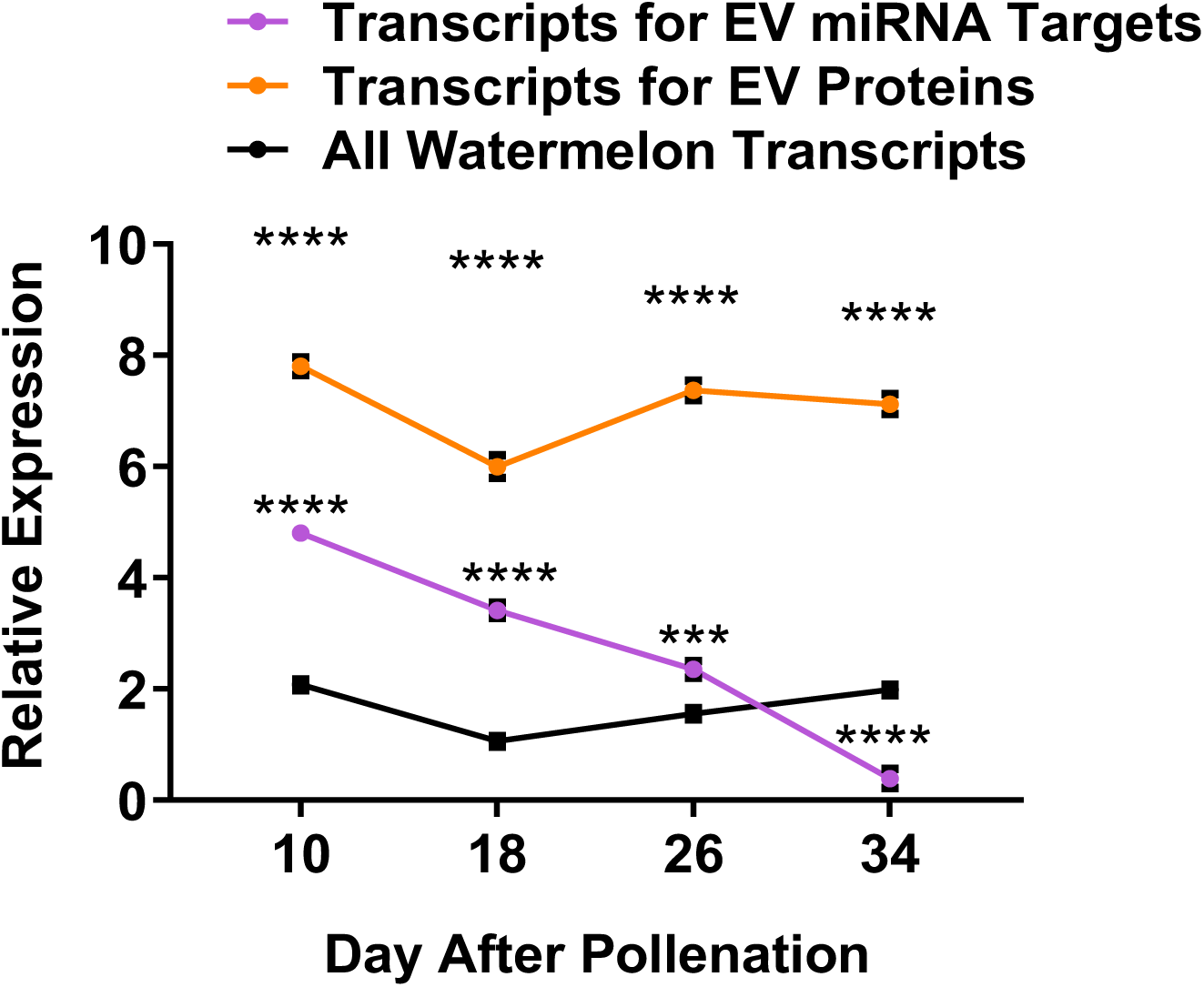
Dynamic changes in watermelon EV miRNA and protein levels across fruit ripening. Levels of miRNA target mRNA and the mRNA for proteins found in watermelon EVs were mapped to mRNA levels across watermelon ripening using a publicly available transcriptomics dataset (Zhu et al., 2017). 105-2944 mRNA per datapoint, two-way ANOVA with Sidak’s multiple comparisons test, comparisons are to ‘all watermelon transcripts’. Data are presented as mean ± SEM.

## Discussion

Here we present the first data on the isolation and characterisation of EVs from watermelon fruit. These EVs are of a size and morphology reminiscent of animal exosomes^53^. We have also reported the average yield of watermelon EVs per millilitre of starting juice and the ratio of particles to DNA, RNA and protein. These characterisations are a first in the plant EV field and are crucial for ensuring that plant EV studies are executed to the same standard as is expected in the mammalian EV field. The particle/protein ratio of 1.85×10^10^ particles/μg protein is considered to be highly pure^54^, suggesting that the methodology presented here is sufficient to isolate EVs from watermelon which are free of protein contamination. However, analysis of other potential contaminant molecules which may be present in watermelon juice, such as glucose and sucrose, is not currently known.

Human exosomes exhibit miRNA^5^ and protein^38^ profiles that differ from their cells of origin due to active loading of cargo, a phenomenon we also observed in watermelon EVs. In watermelon EVs, miRNA sorting appears to be associated with abundance of cellular target proteins; high levels of cytoplasmic mRNA may sequester miRNA away from the site of EV biogenesis, as has been previously identified in humans^40^. However, we acknowledge that protein levels are skewed by post-transcriptional regulation and are therefore an imperfect proxy of mRNA levels. Whilst we did not observe any discriminating sequence or structural motifs between enriched and non-enriched miRNAs in watermelon EVs, characterisation of the full complement of cellular and EV miRNAs is required before this can be confirmed.

Several studies have described EVs or EV-like particles isolated from plants using broadly similar methods to those we employed in this current study^6-15^. Some report vesicles larger than human exosomes and watermelon EVs^8,9,11^, potentially due to vigorous isolation techniques (i.e. prolonged blending) resulting in cellular debris. However, it is also possible that some plant EVs may be constitutively larger, perhaps due to differences in biogenesis. One of the main determinants of the size of vesicles isolated by ultracentrifugation is the final pelleting force^55^. Whilst most studies have pelleted plant EV-like particles at standard forces of 100,000-150,000xg, two studies reported the isolation of EVs at just 40,000xg^6,15^. However, the lack of reporting on the rotors or the pelleting k-factor used to isolate EVs precludes direct comparisons, reinforcing the importance of standardisation in reporting EV isolation and characterisation.

Plastids and MVB are both vesicle-containing organelles^43^ and it is therefore conceivable that both produce plant EVs, perhaps of distinct functionality. More than half of proposed plant EV markers (i.e., those most conserved in EVs between species) are present in chromoplasts, around one third in MVB and one tenth in both, supporting dual origin. Whilst our hypothesis that some watermelon EVs may be generated in plastids is novel, the cell-to-cell movement of plastid components through an unidentified mechanism has been reported previously^56^. Movement of the whole plastid genome between cells was proposed to occur through the movement of intact plastids^56^, but this hypothesis has been criticised due to the large size of plastids relative to the diameter of the plasmodesmata channels between cells^57^. Our data suggest that EVs could be responsible for the transfer of genetic material, as plastid DNA is present at a relatively high abundance in watermelon EVs. DNA sequencing of plant EVs would further confirm this hypothesis.

We have identified a number of potential plant EV marker proteins. We propose that a panel of the potential plant EV markers identified herein – perhaps those with highest fold-enrichment in EVs as compared with cells – be used in future plant EV studies to enable better understanding of the nature of plant EVs as well as ensuring standardisation amongst the field. For example, vesicle-associated protein 4-2 which was identified solely in EVs and is predicted to reside solely in plastids, adenosylhomocysteinase which resides solely in MVB and enolase which is predicted to reside in both plastids and MVB and is also highly abundant in mammalian EVs. Similarly, we recommend that ‘negative’ EV markers from human and animal studies be extended for use in the plant field, as we have shown a number of these to be valid for the characterisation of plant EVs. This includes calnexin and histone 2B.

Previous studies investigating EV-like vesicles in plants have focused on their role in the plant response to infectious agents ^29,58^. Our bioinformatic analyses suggest that plant EVs may also play a role in metabolism and fruit development/ripening. We found that watermelon EVs contain the machinery needed for the metabolism of sucrose and glucose, such as glucose-6-phosphate isomerase and sucrose synthase, suggesting that plant EVs may play a role in the accumulation and metabolism of sugar in fruits, allowing for rapid tissue expansion and making fruit palatable for seed propagation by animals. Growth-regulating factors (GRF) 1, 4, 5 and 9 are predicted targets of several of the most abundant, if not the most enriched, watermelon EV miRNAs, including the miR-396 family and miR-390-3p. GRFs are downregulated during ripening of watermelon and other fruits ^52,59^. A miR-396-GRF network has been implicated in the size, maturation and ripening of several fruits, perhaps through influence on the meristem ^60-62^. Furthermore, miR-159c, which is depleted in watermelon EVs, is also implicated in fruit development and ripening via its targeting of MYB33 ^63,64^; this transcription factor is downregulated during watermelon ripening. Overexpression of miR-159 results in fruit formation in the absence of fertilisation^64,65^ and downregulation of miR-159 results in the production of more spherical fruit in *Arabidopsis thaliana*^66^. Furthermore, three miRNAs enriched (cla-miR319a, miR399g and cla-miR408) and three depleted (cla-miR393, miR167a and cla-miR399a) in watermelon EVs target ethylene-responsive transcription factors and ethylene producing enzymes, suggesting that they may be involved in ethylene-dependent fruit ripening. Together, these data suggest that watermelon EV miRNAs could be involved in the initiation and control of fruit development and ripening.

The potential role of plant EVs in fruit ripening represents a novel finding and could, with further research, aid developing technologies to manipulate this process with the aim of reducing the economic burden of fruit wastage due to premature ripening and subsequent spoiling of fruits before they reach the end consumer. Furthermore, understanding cell-cell communication through EVs in plants could be of use in agricultural grafting, aiding in the production of crops able to develop fruits/vegetables in substandard environments, thereby increasing global food production. Grafted plants share genetic material, with the genetic material from one individual transferring the other ^67-71^. Some have hypothesised that this results from whole nuclei or organelles passing between the individuals following cell disruption during the grafting process (reviewed in ^72^). However, as mentioned above, the small size of the plasmodesmata^73^ communication channels between cells relative to nuclei or organelles questions the feasibility of this occurring ^57^. Therefore, the current prediction of a potential plastid origin for watermelon EVs, especially the plastid DNA they contain, raises the possibility that EVs are a vector for this genetic exchange between grafted species. Further understanding of the role for EVs in grafting could, therefore, aid in increasing crop yield and resilience.

This study contains the first detailed characterisation of plant EVs alongside exploration of the potential functions of watermelon EV cargo. It is hoped that the observations and suggestions made herein, including the panel of plant-specific EV markers, will enable clarity and rigor in the plant EV field moving forward.

## Methods

### Materials

Unless otherwise stated, all reagents were purchased from Sigma-Aldrich.

### Extracellular vesicle isolation

We have submitted all relevant data of our experiments to the EV-TRACK knowledgebase (EV-TRACK ID: EV190074)^74^.Watermelons (n=9) were purchased from a range of local greengrocers and supermarkets. As the variety of watermelons is usually not given in greengrocers or supermarkets, an effort was made to select visually similar fruits. All were of a medium size (i.e. not described as ‘giant’ or ‘baby’) and exhibited dark and light green stripes on their rind. Watermelon were from a range of countries of origin and were purchased throughout the year so as to best reflect the fruits available to consumers. The watermelon mesocarp was removed from the rind with a knife and the weight of dissected mesocarp recorded. Dissected mesocarp was pulsed for 1 second in a standard kitchen blender. This brief blending was sufficient to disrupt the structure of the mesocarp, but did not break open the seeds or produce a homogenate. Large tissue fragments were removed using a coarse sieve to leave an opaque juice. EV isolation was then performed using differential ultracentrifugation at 4 °C with a Sorvall Discovery 100SE centrifuge set to ‘RCFavg’. Firstly, the juice was centrifuged at 1,000xg for 10 minutes using the A-621 fixed-angle rotor. The supernatant was then centrifuged twice at 10,000xg for 10 minutes using A-621 fixed angle rotor to remove the remaining large fragments of tissue, apoptotic bodies and whole organelles such as plastids. The resulting supernatant was passed through a 0.22µm filter (Merck Millipore) then centrifuged at 100,000xg for 90 minutes (k-factor 207.11) in sterile tubes using the T-1250 fixed-angle rotor. The supernatant was discarded, and the pellet was resuspended in 200µl of sterile phosphate buffered saline minus magnesium and calcium. EV size was determined using nanoparticle tracking analysis (NTA) on a NanoSight LM10 (Malvern). Per sample, 3×60 second videos taken and were analysed using the NTA 2.3 software. EV morphology was determined using transmission electron microscopy following negative staining with 1% uranyl acetate. Images were captured on a JEM1400 transmission electron microscope (Jeol) at 120kV using a 1k CCD camera (Advanced Microscopy Techniques, Corp.). Watermelon EVs were compared with those isolated using the same protocol as for watermelon EVs from culture medium conditioned by Caco-2 cells (ECACC #86010202; approximately 1.75 × 10^7^ cells in 50ml culture medium) for 72 hours. Culture medium comprised of Dulbecco’s Modified Eagle Medium (DMEM), 2mM non-essential amino acids, 4mM glutamine, 100U/L penicillin, 100μg/L streptomycin and 10% (v/v) fetal bovine serum (Life Technologies Ltd).

### miRNA analysis

RNA was isolated from watermelon (n=3) EVs or matched mesocarp cells (1,000xg centrifugation pellet, k-factor 50430.28) using the miRVana miRNA Isolation Kit (Thermo Fisher Scientific) with the addition of the Plant RNA Isolation Aid (Thermo Fisher Scientific) according to the manufacturer’s instructions. Following lysis and prior to RNA isolation, samples were spiked with cel-miR-39 (Qiagen) to allow for normalisation of extraction efficiency. Reverse transcription was performed on 125ng of RNA – quantified using the NanoDrop 2000c – using the miScript Plant RT kit (Qiagen). Currently, no products are commercially available for the profiling of watermelon miRNAs. However, the high level of miRNA homology between plant species (discussed in ^75^) allowed profiling of watermelon miRNAs using a rice (*Oryza sativa*) qPCR array, which quantifies 84 miRNAs. cDNA samples from EVs and cells were analysed using the Rice miFinder qPCR array (Qiagen). snoR10, snoR31, snoR5-1a, U15 and U65-2 qPCR array housekeeping genes were constant in cells and EVs and were used alongside the internal reverse transcription control and cel-miR-39 for normalisation. Of the miRNAs on the array, 83 were detected. The melting temperature of each miRNA was compared between qPCR array plates; a difference of 2°C or more suggests that a different sequence had been amplified, such miRNAs were removed from analysis (17 miRNAs). As an additional stringency measure, any rice miRNA with a sequence containing >2 deviations from the published watermelon sequence (Frizzi et al., 2014, Liu et al., 2013) were excluded (38 miRNAs). 28 miRNAs were taken forward for further analysis.

### Western blot

Caco-2 cells and EVs were lysed in 10x radioimmunoprecipitation assay buffer (Millipore), with the concentrated buffer being diluted in distilled water for the cells and in the extracellular vesicle solution for the extracellular vesicles. A total of 30ng of protein was loaded into a 7.5% or 10% polyacrylamide gel (Bio-Rad) in reducing conditions. Following gel electrophoresis, proteins were transferred onto a nitrocellulose membrane (GE Healthcare). Membranes were blocked in 5% bovine serum albumin in tris-buffered saline for 1 hour before being incubated with primary antibodies for ALIX (final concentration 1μg/ml; ab24335; Abcam), CD63 (0.5μg/ml; ab199921; Abcam) or CANX (Calnexin; 1μg/ml; A303-695A; Bethyl Laboratories) overnight at 4°C. Secondary antibody was IRDye 800CW conjugated anti-rabbit IgG (0.2μg/ml; 926-32213; Li-Cor Biosciences). Blots were incubated for 1 hour with secondary antibody and then imaged on the Li-Cor Odyssey Sa (Li-Cor Biosciences).

### Proteomics

Watermelon (n=3) cells and matched extracellular vesicles were lysed in10x radioimmunoprecipitation assay buffer (Millipore), as described above for Caco-2 cells and EVs. A total of 600µg of protein – quantified using the Bradford protein assay (Bio-Rad) – was run 5mm into a 10% polyacrylamide gel (Bio-Rad) prior to fixation and staining using Imperial Protein Stain. The excised protein band was dehydrated using acetonitrile then dried via vacuum centrifugation. The proteins in the dried gel piece were reduced with 10mM dithiothreitol then alkylated with 55mM iodoacetamide. The gel piece was washed twice with 25mM ammonium bicarbonate followed by acetonitrile and vacuum centrifugation. Finally, the proteins in the gel were digested in trypsin overnight at 37°C, producing peptides for mass spectrometry analysis. Digested samples were analysed by liquid chromatography coupled-mass spectrometry/mass spectrometry (LC-MS/MS) using an UltiMate® 3000 Rapid Separation LC (Dionex Corporation) coupled to an Orbitrap Elite (Thermo Fisher Scientific, USA) mass spectrometer. Peptide mixtures were separated for 44 min at 300nl/min-1 using a 1.7µM Ethylene Bridged Hybrid C18 analytical column (75 mm x 250μm internal diameter; Waters) and 0.1% formic acid in a gradient of 8-33% acetonitrile. Detected peptides were then selected for fragmentation automatically by data dependant analysis. The identified peptides were mapped using Mascot 2.5.1 (Matrix Science UK), to the *Citrullus lanatus* proteome. Mascot 2.5.1 was used with a fragment tolerance of 0.60 Da (Monoisotopic), a parent tolerance of 5.0 parts per million (Monoisotopic), fixed modifications of +57 on C (Carbamidomethyl), variable modifications of +16 on M (Oxidation). A maximum of 1 missed cleavage was permitted. Data were validated using Scaffold (Proteome Software, USA), employing the following thresholds: 95% protein identification certainty, 1 for the minimum number of unique peptides mapping to each protein, and 80% peptide identification certainty.

### DNA Analysis

DNA was extracted from watermelon (n=3) cells and matched EVs using the ISOLATE II Plant DNA Kit (Bioline) according the manufacturer’s instructions. Briefly, watermelon cells were ground into a powder using a pestle and mortar with the aid of liquid nitrogen. Both cells and EVs were lysed using Lysis Buffer PA2. Following DNA extraction, primers towards actin-7 (Forward: 5’-ATGTTCACAACCACTGCCGAACG-3’, Reverse: 5’-AATGAGGGATGGCTGGAAAAGAACTT-3’), rbcL (Forward: 5’-ATGAGTTGTAGGGAGGGACTTATGTCACC’3’, Reverse: 5’-GATTTTCTTCTCCAGGAACAGGCTCG-3’) and ccmFn (Forward: 5’-CCGGCCATAGGTTCGAATCCTG-3’, Reverse: 5’-CCGGCCATAGGTTCGAATCCTG-3’) were used to amplify nuclear, plastid and mitochondrial specific DNA respectively from equal amounts of starting DNA using the Fast Cycling PCR Kit (Qiagen). PCR products were run on a 2% agarose gel containing GelRed (Biotium) for band visualisation. Densitometry was conducted using the ImageStudio lite (Li-Cor). The value of any non-specific amplification in negative controls was removed from all densitometry values for the respective primer’s product.

### Bioinformatics analysis

*Oryza sativa* miRNAs were aligned with published watermelon miRNAs using the blastn mode of BLAST, with those miRNAs with <2 mismatches along the length of the miRNA (i.e. <10% deviation from the watermelon sequence) being considered homologous. T-Coffee Multiple Sequence Alignment was used to create a phylogenetic tree of watermelon miRNA sequences. Prediction of miRNA targets was carried out using psRNAtarget ^76^ using the *Citrullus lanatus* (watermelon), transcript, Cucurbit Genomics Database, version 1^77^. FASTA sequences for watermelon proteins were procured from the Cucurbit Genome Database ^77^ and were aligned with those from other plant species and humans using the blastp mode of BLAST. An alignment was considered to predict a homolog if the query cover was at least 70%, with 50% identity within this region. In order to predict the subcellular location of EV biogenesis, the known/annotated subcellular localisation of watermelon EV proteins was obtained from the Cucurbit Genome Database^77^. The presence of motifs required for sorting into different subcellular locations was also used to predict subcellular localisation using the CELLO software^78^. Alignment was also performed to the proteome of watermelon chromoplasts^50^, the late endosome/MVB proteome of *Arabidopsis thaliana* ^51^ and transcriptomics data from ripening watermelon fruit ^52^. *Arabidopsis thaliana* MVB data was used as no watermelon MVB proteome currently exists in the literature. Hierarchical clustering analysis and principal components analysis was performed using the ClustVis R package ^79^. Proteins and predicted miRNA targets were analysed for pathway overrepresentation against the whole watermelon proteome using the Bonferroni test; adjusted p-values of <0.05 were considered significant. This procedure was carried out using the Cucurbit Genome Database.

### Statistical analysis

All statistical analysis was performed on GraphPad Prism 8, unless otherwise stated. All measurements reported are on distinct samples. No samples were measured more than once for the same parameter, except where technical replicates were taken. In the case of technical replicates, the mean of these replicates is reported. The individual statistical tests are given in the relevant methods and figures. In brief, all statistical tests on continuous variables between two groups were analysed using the Mann-Whitney test. When more than two groups were compared over time, a two-way ANOVA with Sidak’s multiple comparisons test was used. Two-sided tests were used for all analyses.

## Data availability

The proteomics and miRNA qPCR array data that support the findings of this study are available in Vesiclepedia (http://microvesicles.org/index.html) with the identifier currently pending. We will provide the identifier as soon as it is available. Furthermore, The authors declare that the proteomics, miRNA qPCR array and miRNA predicted target data supporting the findings of this study are also available within the supplementary information files.

## Acknowledgements

Mass spectrometry was conducted by the University of Manchester Bio-MS core facility. This work was funded through a BBSRC doctoral training partnership PhD project.

## References

1 Brown, L., Wolf, J. M., Prados-Rosales, R. & Casadevall, A. Through the wall: extracellular vesicles in Gram-positive bacteria, mycobacteria and fungi. Nature Reviews Microbiology 13, 620–630, doi:10.1038/nrmicro3480 (2015).

2 van Niel, G., D’Angelo, G. & Raposo, G. Shedding light on the cell biology of extracellular vesicles. Nature Reviews Molecular Cell Biology 19, 213–228, doi:10.1038/nrm.2017.125 (2018).

3 Kosaka, N. et al. Secretory Mechanisms and Intercellular Transfer of MicroRNAs in Living Cells. Journal of Biological Chemistry 285, 17442–17452, doi:10.1074/jbc.M110.107821 (2010).

4 Mittelbrunn, M. et al. Unidirectional transfer of microRNA-loaded exosomes from T cells to antigen-presenting cells. Nature Communications 2, doi:10.1038/ncomms1285 (2011).

5 Valadi, H. et al. Exosome-mediated transfer of mRNAs and microRNAs is a novel mechanism of genetic exchange between cells. Nat Cell Biol 9, 654–659, doi:ncb1596 [pii] 10.1038/ncb1596 (2007).

6 Rutter, B. D. & Innes, R. W. Extracellular Vesicles Isolated from the Leaf Apoplast Carry Stress-Response Proteins. Plant Physiology 173, 728-741, doi:10.1104/pp.16.01253 (2017).

7 Deng, Z. et al. Broccoli-Derived Nanoparticle Inhibits Mouse Colitis by Activating Dendritic Cell AMP-Activated Protein Kinase. Mol Ther 25, 1641–1654, doi:10.1016/j.ymthe.2017.01.025 (2017).

8 Mu, J. Y. et al. Interspecies communication between plant and mouse gut host cells through edible plant derived exosome-like nanoparticles. Molecular Nutrition & Food Research 58, 1561–1573, doi:10.1002/mnfr.201300729 (2014).

9 Zhang, M. et al. Edible ginger-derived nanoparticles: A novel therapeutic approach for the prevention and treatment of inflammatory bowel disease and colitis-associated cancer. Biomaterials 101, 321–340, doi:10.1016/j.biomaterials.2016.06.018 (2016).

10 Zhang, M., Wang, X., Han, M. K., Collins, J. F. & Merlin, D. Oral administration of ginger-derived nanolipids loaded with siRNA as a novel approach for efficient siRNA drug delivery to treat ulcerative colitis. Nanomedicine (Lond) 12, 1927–1943, doi:10.2217/nnm-2017-0196 (2017).

11 Ju, S. W. et al. Grape Exosome-like Nanoparticles Induce Intestinal Stem Cells and Protect Mice From DSS-Induced Colitis. Molecular Therapy 21, 1345–1357, doi:10.1038/mt.2013.64 (2013).

12 Pérez-Bermúdez, P., Blesa, J., Soriano, J. M. & Marcilla, A. Extracellular vesicles in food: Experimental evidence of their secretion in grape fruits. Eur J Pharm Sci 98, 40–50, doi:10.1016/j.ejps.2016.09.022 (2017).

13 Wang, B. M. et al. Targeted Drug Delivery to Intestinal Macrophages by Bioactive Nanovesicles Released from Grapefruit. Molecular Therapy 22, 522–534, doi:10.1038/mt.2013.190 (2014).

14 Raimondo, S. et al. Citrus limon-derived nanovesicles inhibit cancer cell proliferation and suppress CML xenograft growth by inducing TRAIL-mediated cell death. Oncotarget 6, 19514–19527, doi:10.18632/oncotarget.4004 (2015).

15 Regente, M. et al. Vesicular fractions of sunflower apoplastic fluids are associated with potential exosome marker proteins. Febs Letters 583, 3363–3366, doi:10.1016/j.febslet.2009.09.041 (2009).

16 Xiao, J. et al. Identification of exosome-like nanoparticle-derived microRNAs from 11 edible fruits and vegetables. PeerJ 6, e5186, doi:10.7717/peerj.5186 (2018).

17 Raposo, G. et al. B lymphocytes secrete antigen-presenting vesicles. Journal of Experimental Medicine 183, 1161–1172, doi:10.1084/jem.183.3.1161 (1996).

18 Futter, C. E., Pearse, A., Hewlett, L. J. & Hopkins, C. R. Multivesicular endosomes containing internalized EGF-EGF receptor complexes mature and then fuse directly with lysosomes. Journal of Cell Biology 132, 1011–1023, doi:10.1083/jcb.132.6.1011 (1996).

19 Donker, R. B. et al. The expression profile of C19MC microRNAs in primary human trophoblast cells and exosomes. Molecular Human Reproduction 18, 417–424, doi:10.1093/molehr/gas013 (2012).

20 Van Niel, G. et al. Intestinal epithelial cells secrete exosome–like vesicles. Gastroenterology 121, 337–349, doi:http://dx.doi.org/10.1053/gast.2001.26263 (2001).

21 Hunter, M. P. et al. Detection of microRNA Expression in Human Peripheral Blood Microvesicles. Plos One 3, 11, doi:10.1371/journal.pone.0003694 (2008).

22 Heijnen, H. F. G., Schiel, A. E., Fijnheer, R., Geuze, H. J. & Sixma, J. J. Activated platelets release two types of membrane vesicles: Microvesicles by surface shedding and exosomes derived from exocytosis of multivesicular bodies and alpha-granules. Blood 94, 3791–3799 (1999).

23 Villarroya-Beltri, C. et al. Sumoylated hnRNPA2B1 controls the sorting of miRNAs into exosomes through binding to specific motifs. Nature Communications 4, 10, doi:10.1038/ncomms3980 (2013).

24 Moreno-Gonzalo, O., Fernandez-Delgado, I. & Sanchez-Madrid, F. Post-translational add-ons mark the path in exosomal protein sorting. Cellular and Molecular Life Sciences 75, 1–19, doi:10.1007/s00018-017-2690-y (2018).

25 Holme, P. A., Brosstad, F. & Solum, N. O. The difference between platelet and plasma FXIII used to study the mechanism of platelet microvesicle formation. Thrombosis and Haemostasis 70, 681–686 (1993).

26 Hristov, M., Erl, W., Linder, S. & Weber, P. C. Apoptotic bodies from endothelial cells enhance the number and initiate the differentiation of human endothelial progenitor cells in vitro. Blood 104, 2761–2766 (2004).

27 Cotter, T. G., Lennon, S. V., Glynn, J. M. & Green, D. R. Microfilament-disrupting agents prevent the formation of apoptotic bodies in tumour cells undergoing apoptosis. Cancer Research 52, 997–1005 (1992).

28 Jiang, L. et al. Determining the contents and cell origins of apoptotic bodies by flow cytometry. Scientific Reports 7, doi:10.1038/s41598-017-14305-z (2017).

29 An, Q. L., Huckelhoven, R., Kogel, K. H. & Van Bel, A. J. E. Multivesicular bodies participate in a cell wall-associated defence response in barley leaves attacked by the pathogenic powdery mildew fungus. Cellular Microbiology 8, 1009–1019, doi:10.1111/j.1462-5822.2006.00683.x (2006).

30 Baldrich, P. et al. Plant Extracellular Vesicles Contain Diverse Small RNA Species and Are Enriched in 10-to 17-Nucleotide “Tiny” RNAs. Plant Cell 31, 315–324, doi:10.1105/tpc.18.00872 (2019).

31 Cai, Q. et al. Plants send small RNAs in extracellular vesicles to fungal pathogen to silence virulence genes. Science 360, 1126–1129, doi:10.1126/science.aar4142 (2018).

32 Rutter, B. D. & Innes, R. W. Extracellular vesicles as key mediators of plant-microbe interactions. Current Opinion in Plant Biology 44, 16–22, doi:10.1016/j.pbi.2018.01.008 (2018).

33 Regente, M., Monzon, G. C. & de la Canal, L. Phospholipids are present in extracellular fluids of imbibing sunflower seeds and are modulated by hormonal treatments. Journal of Experimental Botany 59, 553–562, doi:10.1093/jxb/erm329 (2008).

34 Gonorazky, G., Laxalt, A. M., Testerink, C., Munnik, T. & De La Canal, L. Phosphatidylinositol 4-phosphate accumulates extracellularly upon xylanase treatment in tomato cell suspensions. Plant Cell and Environment 31, 1051–1062, doi:10.1111/j.1365-3040.2008.01818.x (2008).

35 Regente, M., Pinedo, M., Elizalde, M. & de la Canal, L. Apoplastic exosome-like vesicles A new way of protein secretion in plants? Plant Signaling & Behavior 7, 544–546, doi:10.4161/psb.19675 (2012).

36 Baietti, M. F. et al. Syndecan-syntenin-ALIX regulates the biogenesis of exosomes. Nature Cell Biology 14, 677–685, doi:10.1038/ncb2502 (2012).

37 Iavello, A. et al. Role of Alix in miRNA packaging during extracellular vesicle biogenesis. International Journal of Molecular Medicine 37, 958–966, doi:10.3892/ijmm.2016.2488 (2016).

38 Haraszti, R. A. et al. High-resolution proteomic and lipidomic analysis of exosomes and microvesicles from different cell sources. Journal of Extracellular Vesicles 5, doi:10.3402/jev.v5.32570 (2016).

39 Théry, C. et al. Minimal information for studies of extracellular vesicles 2018 (MISEV2018): a position statement of the International Society for Extracellular Vesicles and update of the MISEV2014 guidelines. Journal of Extracellular Vesicles 8, 1535750, doi:10.1080/20013078.2018.1535750 (2019).

40 Squadrito, M. L. et al. Endogenous RNAs Modulate MicroRNA Sorting to Exosomes and Transfer to Acceptor Cells. Cell Reports 8, 1432–1446, doi:10.1016/j.celrep.2014.07.035 (2014).

41 Kim, D. K. et al. EVpedia: a community web portal for extracellular vesicles research. Bioinformatics 31, 933–939, doi:10.1093/bioinformatics/btu741 (2015).

42 Kalra, H. et al. Vesiclepedia: A Compendium for Extracellular Vesicles with Continuous Community Annotation. Plos Biology 10, doi:10.1371/journal.pbio.1001450 (2012).

43 Lindquist, E., Solymosi, K. & Aronsson, H. Vesicles Are Persistent Features of Different Plastids. Traffic 17, 1125–1138, doi:10.1111/tra.12427 (2016).

44 Westphal, S., Soll, J. & Vothknecht, U. C. A vesicle transport system inside chloroplasts. Febs Letters 506, 257–261, doi:10.1016/s0014-5793(01)02931-3 (2001).

45 Morre, D. J., Sellden, G., Sundqvist, C. & Sandelius, A. S. STROMAL LOW-TEMPERATURE COMPARTMENT DERIVED FROM THE INNER MEMBRANE OF THE CHLOROPLAST ENVELOPE. Plant Physiology 97, 1558–1564, doi:10.1104/pp.97.4.1558 (1991).

46 Zhang, L., Kato, Y., Otters, S., Vothknecht, U. C. & Sakamoto, W. Essential Role of VIPP1 in Chloroplast Envelope Maintenance in Arabidopsis. Plant Cell 24, 3695–3707, doi:10.1105/tpc.112.103606 (2012).

47 Aseeva, E. et al. Vipp1 is required for basic thylakoid membrane formation but not for the assembly of thylakoid protein complexes. Plant Physiology and Biochemistry 45, 119–128, doi:10.1016/j.plaphy.2007.01.005 (2007).

48 Kroll, D. et al. VIPP1, a nuclear gene of Arabidopsis thaliana essential for thylakoid membrane formation. Proceedings of the National Academy of Sciences of the United States of America 98, 4238–4242, doi:10.1073/pnas.061500998 (2001).

49 Caiola, M. G. & Canini, A. Ultrastructure of chromoplasts and other plastids in Crocus sativus L. (Iridaceae). Plant Biosystems 138, 43–52, doi:10.1080/11263500410001684116 (2004).

50 Wang, Y. Q. et al. Proteomic analysis of chromoplasts from six crop species reveals insights into chromoplast function and development. Journal of Experimental Botany 64, 949–961, doi:10.1093/jxb/ers375 (2013).

51 Heard, W., Sklenar, J., Tome, D. F. A., Robatzek, S. & Jones, A. M. E. Identification of Regulatory and Cargo Proteins of Endosomal and Secretory Pathways in Arabidopsis thaliana by Proteomic Dissection. Molecular & Cellular Proteomics 14, 1796–1813, doi:10.1074/mcp.M115.050286 (2015).

52 Zhu, Q. L. et al. Comparative transcriptome analysis of two contrasting watermelon genotypes during fruit development and ripening. Bmc Genomics 18, doi:10.1186/s12864-016-3442-3 (2017).

53 Thery, C., Amigorena, S., Raposo, G. & Clayton, A. Isolation and characterization of exosomes from cell culture supernatants and biological fluids. Current protocols in cell biology Chapter 3, Unit 3.22-Unit 23.22, doi:10.1002/0471143030.cb0322s30 (2006).

54 Webber, J. & Clayton, A. How pure are your vesicles? Journal of Extracellular Vesicles 2, 19861, doi:10.3402/jev.v2i0.19861 (2013).

55 Witwer, K. W. et al. Standardization of sample collection, isolation and analysis methods in extracellular vesicle research. Journal of extracellular vesicles 2, doi:10.3402/jev.v2i0.20360 (2013).

56 Thyssen, G., Svab, Z. & Maliga, P. Cell-to-cell movement of plastids in plants. Proceedings of the National Academy of Sciences of the United States of America 109, 2439–2443, doi:10.1073/pnas.1114297109 (2012).

57 Bobik, K. & Burch-Smith, T. M. Chloroplast signaling within, between and beyond cells. Frontiers in Plant Science 6, doi:10.3389/fpls.2015.00781 (2015).

58 Nielsen, M. E., Feechan, A., Boehlenius, H., Ueda, T. & Thordal-Christensen, H. Arabidopsis ARF-GTP exchange factor, GNOM, mediates transport required for innate immunity and focal accumulation of syntaxin PEN1. Proceedings of the National Academy of Sciences of the United States of America 109, 11443–11448, doi:10.1073/pnas.1117596109 (2012).

59 Liu, X. et al. Identification and transcript profiles of citrus growth-regulating factor genes involved in the regulation of leaf and fruit development. Molecular Biology Reports 43, 1059–1067, doi:10.1007/s11033-016-4048-1 (2016).

60 Wang, L. et al. miR396-targeted AtGRF transcription factors are required for coordination of cell division and differentiation during leaf development in Arabidopsis. Journal of Experimental Botany 62, 761–773, doi:10.1093/jxb/erq307 (2011).

61 Wang, L., Du, H. Y. & Wuyun, T. N. Genome-Wide Identification of MicroRNAs and Their Targets in the Leaves and Fruits of Eucommia ulmoides Using High-Throughput Sequencing. Frontiers in Plant Science 7, 15, doi:10.3389/fpls.2016.01632 (2016).

62 Cao, D. Y. et al. Regulations on growth and development in tomato cotyledon, flower and fruit via destruction of miR396 with short tandem target mimic. Plant Science 247, 1–12, doi:10.1016/j.plantsci.2016.02.012 (2016).

63 Csukasi, F. et al. Two strawberry miR159 family members display developmental-specific expression patterns in the fruit receptacle and cooperatively regulate Fa-GAMYB. New Phytologist 195, 47–57, doi:10.1111/j.1469-8137.2012.04134.x (2012).

64 da Silva, E. M. et al. microRNA159-targeted SIGAMYB transcription factors are required for fruit set in tomato. Plant Journal 92, 95–109, doi:10.1111/tpj.13637 (2017).

65 Palatnik, J. F. et al. Sequence and expression differences underlie functional specialization of Arabidopsis microRNAs miR159 and miR319. Dev Cell 13, 115–125, doi:10.1016/j.devcel.2007.04.012 (2007).

66 Allen, R. S. et al. Genetic analysis reveals functional redundancy and the major target genes of the Arabidopsis miR159 family. Proceedings of the National Academy of Sciences of the United States of America 104, 16371–16376, doi:10.1073/pnas.0707653104 (2007).

67 Stegemann, S. & Bock, R. Exchange of Genetic Material Between Cells in Plant Tissue Grafts. Science 324, 649–651, doi:10.1126/science.1170397 (2009).

68 Molnar, A. et al. Small Silencing RNAs in Plants Are Mobile and Direct Epigenetic Modification in Recipient Cells. Science 328, 872–875, doi:10.1126/science.1187959 (2010).

69 Fragoso, V., Goddard, H., Baldwin, I. T. & Kim, S. G. A simple and efficient micrografting method for stably transformed Nicotiana attenuata plants to examine shoot-root signaling. Plant Methods 7, 8, doi:10.1186/1746-4811-7-34 (2011).

70 Melnyk, C. W., Molnar, A., Bassett, A. & Baulcombe, D. C. Mobile 24 nt Small RNAs Direct Transcriptional Gene Silencing in the Root Meristems of Arabidopsis thaliana. Current Biology 21, 1678–1683, doi:10.1016/j.cub.2011.08.065 (2011).

71 Buhtz, A., Pieritz, J., Springer, F. & Kehr, J. Phloem small RNAs, nutrient stress responses, and systemic mobility. Bmc Plant Biology 10, 13, doi:10.1186/1471-2229-10-64 (2010).

72 Wang, J., Jiang, L. B. & Wu, R. L. Plant grafting: how genetic exchange promotes vascular reconnection. New Phytologist 214, 56–65, doi:10.1111/nph.14383 (2017).

73 Pina, A., Errea, P. & Martens, H. J. Graft union formation and cell-to-cell communication via plasmodesmata in compatible and incompatible stem unions of Prunus spp. Scientia Horticulturae 143, 144–150, doi:10.1016/j.scienta.2012.06.017 (2012).

74 Van Deun, J. et al. EV-TRACK: transparent reporting and centralizing knowledge in extracellular vesicle research. Nature Methods 14, 228–232, doi:10.1038/nmeth.4185 (2017).

75 Li, A. L. & Mao, L. Evolution of plant microRNA gene families. Cell Research 17, 212–218, doi:10.1038/sj.cr.7310113 (2007).

76 Dai, X. B. & Zhao, P. X. psRNATarget: a plant small RNA target analysis server. Nucleic Acids Research 39, W155–W159, doi:10.1093/nar/gkr319 (2011).

77 Feilab. ‘Cucurbit Genome Database’, <http://cucurbitgenomics.org/> (2017).

78 Yu, C.-S., Chen, Y.-C., Lu, C.-H. & Hwang, J.-K. Prediction of protein subcellular localization. Proteins: Structure, Function, and Bioinformatics 64, 643–651, doi:10.1002/prot.21018 (2006).

79 Metsalu, T. & Vilo, J. ClustVis: a web tool for visualizing clustering of multivariate data using Principal Component Analysis and heatmap. Nucleic Acids Research 43, W566–W570, doi:10.1093/nar/gkv468 (2015).

